# Kir7.1 is the physiological target for hormones and steroids that regulate uteroplacental function

**DOI:** 10.1101/2023.04.11.536476

**Authors:** Monika Haoui, Citlalli Vergara, Polina V. Lishko

**Affiliations:** Department of Molecular and Cell Biology, University of California, Berkeley, CA; Center for Reproductive Longevity and Equality at Buck Institute for Research on Aging, Novato, CA; Department of Cell Biology and Physiology, Washington University in St. Louis, School of Medicine, St. Louis, MO

**Keywords:** K_ir_7.1, myometrium, placenta, fetoplacental unit, labyrinth, preeclampsia, parturition, steroid hormones, progesterone, preterm labor

## Abstract

Preterm birth is a multifactorial syndrome that is detrimental to the well-being of both the mother and the newborn. During normal gestation, the myometrium is maintained in a quiescent state by the action of progesterone. As a steroid hormone, progesterone is thought to modify uterine and placental morphology by altering gene expression, but a nongenomic mode of action has long been suspected. Here we reveal that progesterone activates both human and murine inwardly rectifying potassium channel Kir7.1, which is expressed in mammalian myometrial smooth muscle and placental pericytes during late gestation. Kir7.1 is also activated by compounds used to prevent premature labor, including the progestogens 17-alpha-hydroxyprogesterone caproate and dydrogesterone, revealing an unexpected mode of action for these drugs. Our results reveal that Kir7.1 is the molecular target of a number of endogenous and synthetic steroids that control uterine excitability and placental function, and is therefore a promising therapeutic target to control utero-placental physiology and support healthy pregnancy.

## Introduction

The uterus is the strongest smooth muscle in the body per mass of tissue, but it must remain quiescent during pregnancy to house and support the developing fetus. As parturition approaches, the uterus undergoes profound physiological changes to allow strong, synchronized myometrial contractions for a successful delivery. At the same time, the placenta, a heavily vascularized organ that supports embryogenesis, also undergoes significant remodeling.

Specifically, the placenta develops a heavily vascularized structure called the labyrinth, which provides nutrient exchange between mother and fetus (1) and in humans produces the majority of progesterone (P4) (2). For more than 50 years, P4 has been recognized as the main factor supporting pregnancy and uterine quiescence, but its mechanism of action on the myometrium is poorly understood (3-6). In most mammals, a fall in P4 levels is required for the onset of labor.

However, P4 is present during parturition in humans and most primates, suggestive of a “functional withdrawal” mechanism (5,7). It is thought that activity changes in the nuclear progesterone receptor underlie functional withdrawal at parturition (5,6,8), but this cannot explain the rapid and reversible relaxation observed when P4 is applied to isolated myometrial strips (3,5). Instead, there is likely to be a membrane and nongenomic target of progesterone that modifies cellular excitability, such as ion channels.

As in all smooth muscle, uterine contractions are initiated by the concerted action of voltage- and ligand-gated ion channels. Excitation is initiated when Ca^2+^ ions enter myometrial cells through voltage-gated Ca^2+^ channels, activating actin-myosin complexes and initiating the cross-bridge cycle (9). K^+^ channels, on the other hand, play a vital role in maintaining quiescence by hyperpolarizing the myolemma, thereby inactivating voltage-gated Ca^2+^ channels. Kir7.1 is one such potassium channel (10-12), whose inhibition and knockdown increases uterine contractility, while reintroducing Kir7.1 after knockdown restores uterine quiescence (13,14).

Interestingly, a decrease in Kir7.1 mRNA expression in the pregnant myometrium coincides with the onset of labor (14).

Here we show that myometrial murine Kir7.1 is expressed in a specific uterine compartment adjacent to the placenta and, moreover, in the placental labyrinth. The labyrinth contains trophoblasts and endothelial cells that are covered with specific mural cells called placental pericytes. The latter control the blood supply to the fetus (1), but the molecular mechanism of this process is poorly understood. Our results reveal that Kir7.1 expression in the murine placenta is specific to placental pericytes and that both human and murine Kir7.1 are activated by progesterone. Moreover, we show that two therapeutic compounds used to prevent preterm labor and potentially treat preeclampsia (15,16) are specific and potent activators of human Kir7.1. Further electrophysiological studies revealed that additional pregnancy-related hormones, including estrogen and dehydroepiandrosterone (DHEA), are key regulators of Kir7.1. Moreover, mifepristone (RU-486), an abortifacient used during early pregnancy, is a Kir7.1 antagonist that opposes the effect of P4. These data thus reveal the previously unknown mechanism of action of mifepristone as well as therapeutic compounds used to maintain pregnancy such as 17-alpha-hydroxyprogesterone caproate and dydrogesterone. Our results provide strong evidence for a nongenomic mechanism behind the rapid control of uterine excitability by progesterone and support the notion that Kir7.1 is a promising molecular target to control uteroplacental physiology.

## Results

### Kir 7.1 is expressed in myometrial smooth muscle and placental pericytes during gestation

The uterus consists of two muscular layers of smooth muscles, a longitudinal outer layer and a circular inner layer (Fig. 1A-E). The circular inner layer surrounds the endometrium, while the longitudinal layer aligns along the long axis of the uterus. While these two layers have been shown to exhibit different contractile characteristics and protein expression profiles, their unique roles in pregnancy have not been established (17-18). Kir7.1 was found to be expressed in both circular and longitudinal myometrial cells surrounding the placenta (Fig. 1D-E) but in neither layer of the myometrial tissue found between placentas in the uterus (Fig. 1B), putting Kir7.1 in close proximity to the main producer of progesterone during pregnancy in humans, the placenta. Pregnancy-related myometrial Kir7.1 expression corroborates previous studies (13); however, its specific localization to the placental side of the uterus was unexpected.

**Figure 1.**
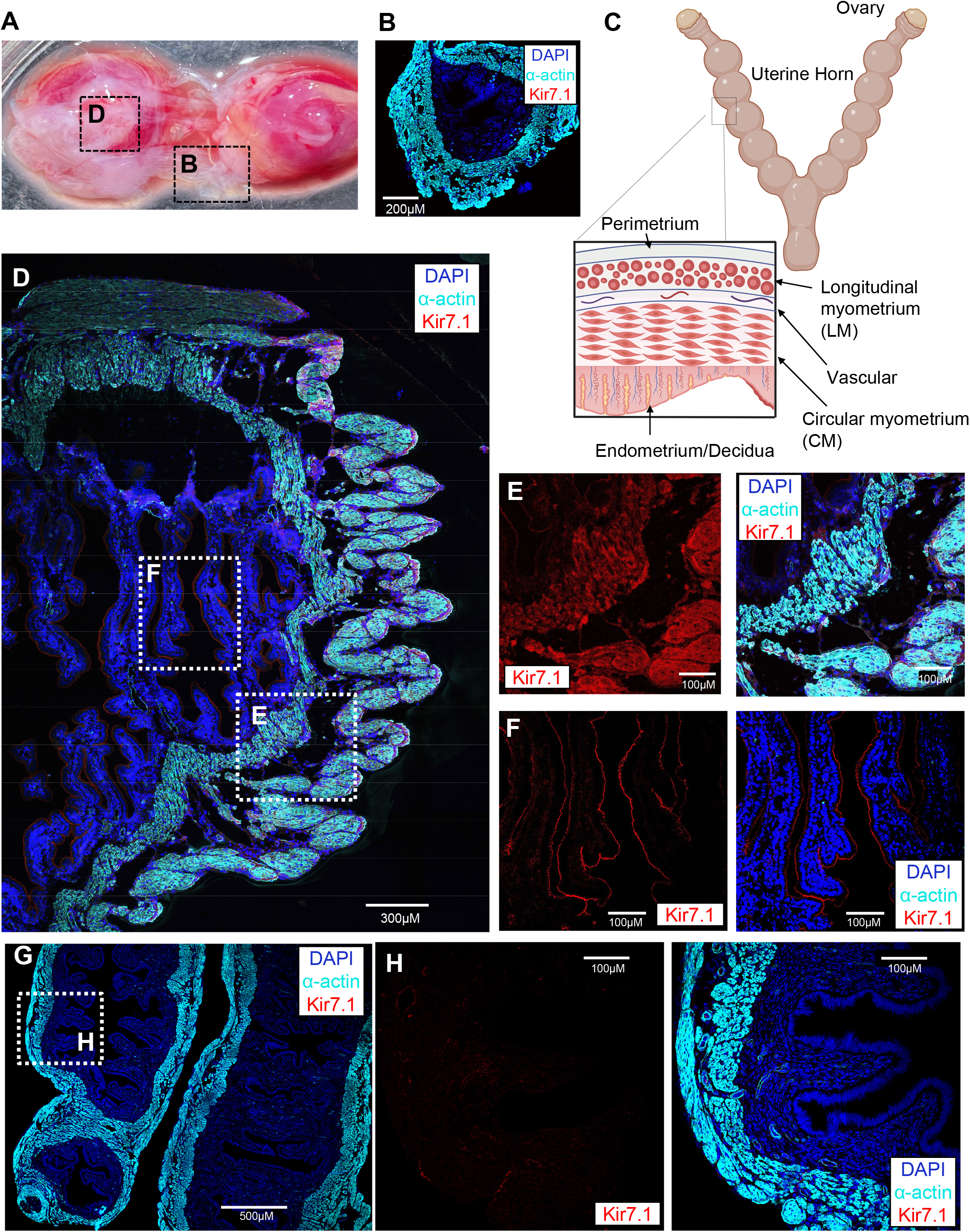
Kir7.1 expression and localization in the murine myometrium. **A**. Isolated mouse uterus at 15.5 days post coitum (dpc) with placentas (red tissue) and myometrium (white tissue) shown. **B**. Immunohistochemistry of myometrial sections between placentas visualized by smooth muscle alpha-actin (cyan). Myocytes from this section lack Kir7.1 expression (red). Cell nuclei are stained by DAPI (blue). **C**. Schematic representation of mouse uterine morphology during gestation. The insert shows different uterine layers, including the endometrium, circular myometrial inner layer, and vascular and longitudinal myometrial layers. Created with BioRender.com. **D**. Immunohistochemistry of the myometrial section at 15.5 dpc adjacent to the placenta shows Kir7.1 expression in both circular and longitudinal myometrial cells (**E**, red), as well as apical endometrium surrounding the placenta (**F**, red). Right panels show merged alpha-actin (cyan), Kir7.1 (red) and DAPI (blue) staining. **G**. Nonpregnant uterus lacks Kir7. **H**. Zoomed-in portion of nonpregnant uterus from (g) with absent Kir7.1 and weak background staining. The right panel shows merged alpha-actin (cyan), Kir7.1 (red) and DAPI (blue) staining.

Recently, we reported that murine Kir7.1 expressed in the choroid plexus and retinal pigment epithelia is potently and specifically activated by progesterone (19, 20) and that such activation is independent of G-protein regulation (20). This finding led us to compare the properties of human and murine Kir7.1 to verify whether human Kir7.1 can be regulated by progesterone in the similar manner. Additionally, the function of this potassium channel and its potential regulation by steroids in the myometrium and placenta were explored. First, we confirmed that Kir7.1 was not expressed in the murine nonpregnant myometrium (Fig. 1G-H), which was supported by both immunohistochemistry studies and by functional interrogation of isolated myometrial cells (Suppl. Fig. 1A-E). In addition to uterine expression, Kir7.1 expression was particularly intense in the murine placental labyrinth (Fig. 2A-D). Vascular myocytes that surround labyrinthic blood vessels (Fig. 2C, F-G) are surrounded by trophoblasts and tightly connect with pericytes (Suppl. Fig. 2). Placental pericytes can be easily distinguished by their characteristic presence of smooth muscle actin (sm-alpha-actin (21), Fig. 2), in contrast to other vascular pericytes that lack sm-alpha-actin (22). The isolated pericytes revealed asymmetrically clustered Kir7.1 expression (Suppl. Fig. 2A-C). Interestingly, Kir7.1 expression was restricted to placental pericytes, select myometrial smooth muscle cells, and apical decidua (Fig. 1-2). At the same time, vascular myocytes that surround placental arterioles and veins and are also visualized by sm-alpha-actin lacked Kir7.1 expression (Fig. 2F-G and Suppl. Fig. 2D). However, Kir7.1 was detected in the membrane junctions between pericytes and vascular myocytes, resembling intimate pericytial-muscular contacts in the brain microvasculature (Suppl. Fig 2D; (22)).

**Figure 2.**
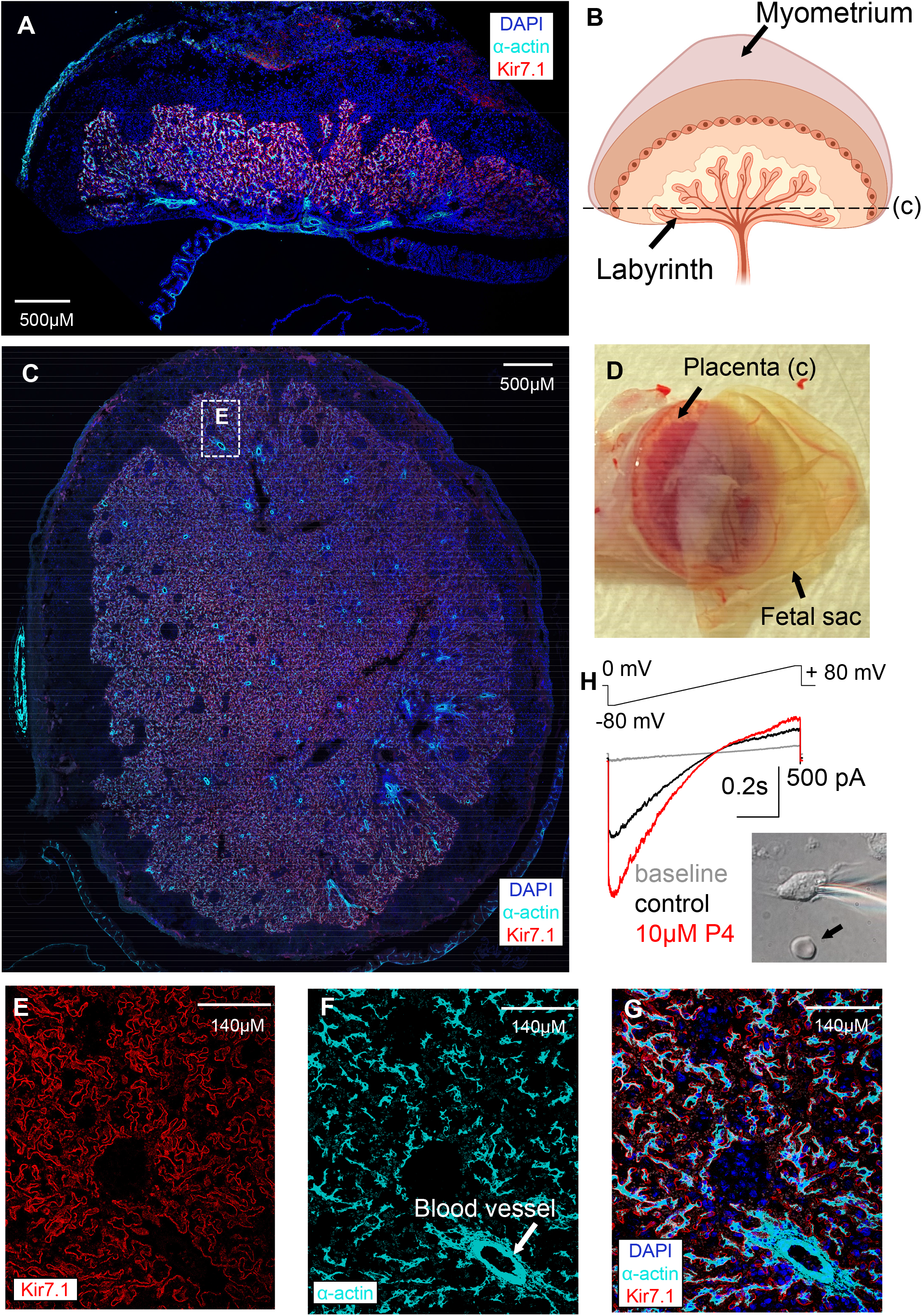
Kir7.1 expression and localization in the murine placental labyrinth. **A**. Immunostaining of the isolated mouse utero-placental unit at 15.5 dpc. The coronal section through the unit revealed intense Kir.7.1 expression (red) and smooth muscle alpha-actin (cyan) staining in the labyrinth. Cell nuclei are stained by DAPI (blue). **B**. Schematic of the utero-placental unit with labyrinth and the horizontal section plain through it (c). Created with BioRender.com. **C**. Horizontal cross-section through labyrinth at 15.5 dpc. Kir7.1 (red) is abundant in placental pericytes visualized by characteristic smooth muscle alpha-actin (cyan) staining. **D**. Isolated mouse utero-placental unit. **E**. Zoomed in section of (c) with individual Kir7.1 staining (red) shows clear membrane expression of the channel. **F**. Zoomed-in section of (c) with individual smooth muscle alpha-actin (cyan) staining shows pericytes and smooth muscle vasculature surrounding blood vessels (indicated by arrow). **G**. Same panels as (e) and (f) merged. Kir7.1 expression is absent from blood vessel vasculature. Cell nuclei are stained by DAPI (blue). **H**. Representative traces recorded from isolated placental pericytes (DIC image of the cell is shown in the insert) show characteristic inward rectification that is further stimulated by 10 µM progesterone (P4).

### Kir 7.1 is activated by progesterone in the pregnant myometrium

The functional presence of Kir7.1 was further confirmed by direct electrophysiological recordings from isolated murine myometrial cells (Supp. Fig 1F-I) and isolated murine placental pericytes (Fig. 2H and Suppl. Fig. 3) from five 15.5 days post coitum (dpc) mice. At the same time, isolated myocytes from nonpregnant murine uteri lacked any Kir7.1 activity (Suppl. Fig. 1A-C). Individual isolation of myocytes and pericytes for electrophysiological interrogation requires mild proteolytic digestion of the murine utero-placental unit. This procedure leads to partial internalization of Kir7.1 and, hence, its low surface representation, which resulted in significant variabilities between Kir7.1 current densities in both isolated myocytes and pericytes (Suppl. Fig 1F-I and Suppl. 3A-C). To circumvent this, we explored murine and human Kir7.1 sensitivity to steroid compounds by using a heterologous expression system and overexpressed Kir7.1 in either HEK293 cells or cultured human myometrial cells (Suppl. Fig. 4A-H and Suppl. Fig. 5A-B).

### Progesterone activates both human and murine Kir 7.1

It was determined that both murine and human Kir7.1 were similarly potentiated by P4 (Suppl. Fig. 4I-J). Kir7.1 is an evolutionarily conserved ion channel with 92.6% identity between humans and rodents (mouse and rat). Therefore, due to this sequence homology, it is expected for Kir7.1 to exert the same biological function in murine and human tissues. The EC_50_ of P4 potentiation of Kir7.1 was comparable between murine Kir7.1 expressed in HEK293 cells (Suppl. Fig. 5C) and human Kir7.1 expressed in either HEK293 or myometrium and were between 9 and 11.5 μM (Suppl. Fig. 4I). This is in line with physiological concentrations that progesterone reached during pregnancy. Since P4 is produced by the uterus and placenta directly, with the human placenta producing 250 mg/day P4 (23), it has been shown that local steroid concentrations are significantly greater than the concentration of these steroids in the blood plasma (24), at least in humans. Therefore, endogenous P4 is not only locally produced but is available at physiological concentrations to activate Kir7.1.

### Human Kir 7.1 responds to a number of endogenous steroids

While P4 plays a dominant role in maintaining quiescence during pregnancy to accommodate the developing fetus, there are other sex steroid hormones produced during pregnancy that play unique, yet poorly understood, roles. Dehydroepiandrosterone (DHEA) and its sulfated form of DHEA sulfate (DHEA-S) are produced by fetal and maternal adrenals in large amounts for placental estrogen production (25). In the placenta, estradiol (E2) is the most abundant estrogen synthesized. While few synthetic inhibitors of human Kir7.1 are known, endogenous activators have yet to be identified. Therefore, the modulation of human Kir7.1 by pregnancy-related steroids has been explored. The average current fold increase of human Kir7.1 recorded from HEK293 cells that were exposed to 10 µM estrogen, estrone, DHEA, DHEA-S, and cortisol was compared to the potentiation caused by P4 (Fig. 3A-E and Suppl. Fig. 6). Interestingly, among these steroids, only DHEA showed agonistic activity, while its membrane impermeable sulfated analog, DHEA-S, was inactive (Fig. 3A-E and Suppl. Fig. 6). This hinted at the possibility that Kir7.1 is activated by steroids from the intracellular side, for which membrane penetration is necessary.

**Figure 3.**
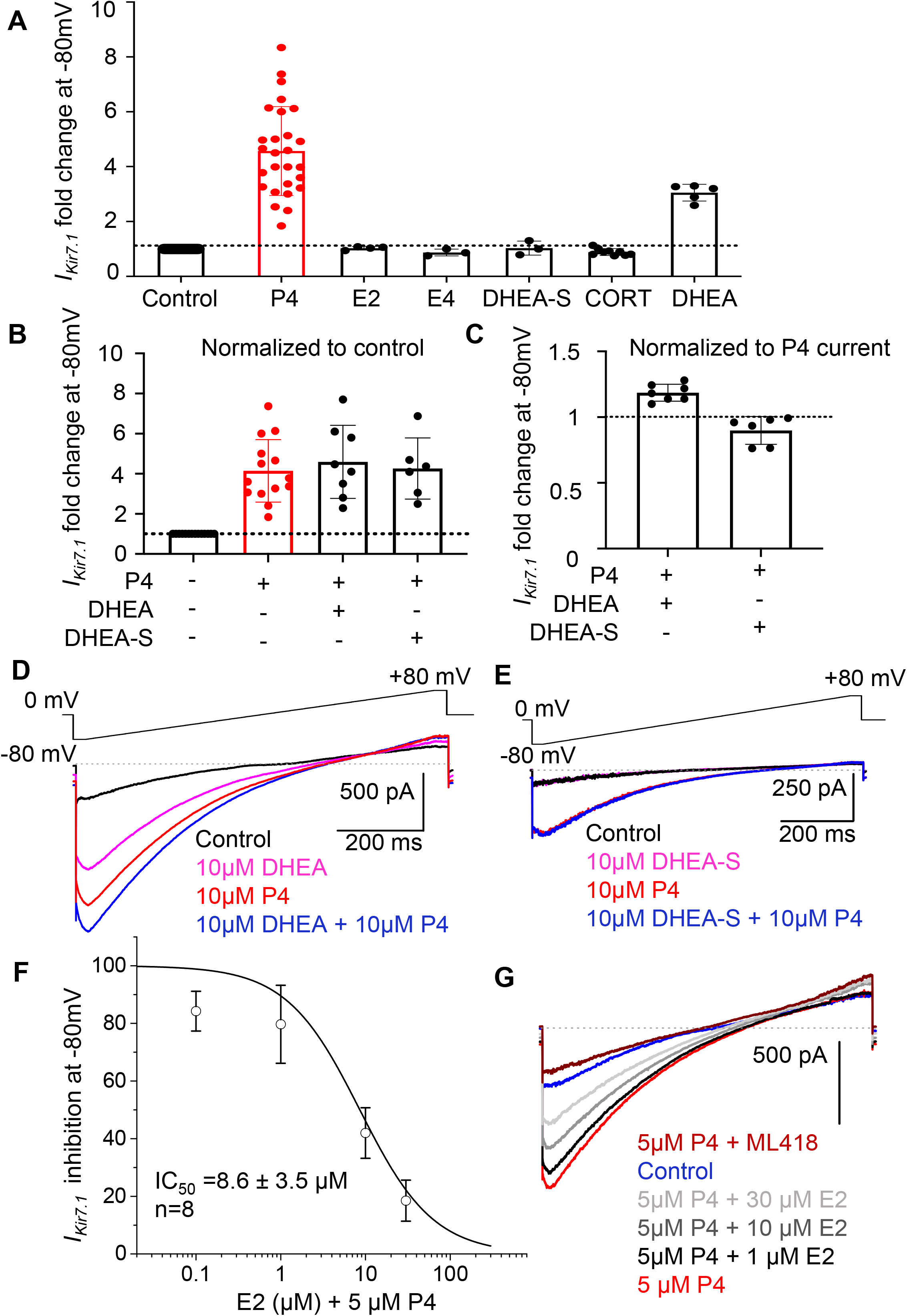
Potentiation of human Kir7.1 by endogenous steroids. **A**. The average current fold increase of human Kir7.1 at -80 mV recorded from recombinant channel expressed in HEK293 cells that were exposed to 10 µM of progesterone (P4), estrogen (E2), estrone (E4), DHEA-S, cortisol (Cort), and DHEA. **B**. Individual and combined application of P4 and DHEA or DHEA-S. Individual steroids were applied in different orders for each cell, followed by P4 combined with DHEA or DHEAS. Data are means ± SEM. **C**. Current densities of combined P4/DHEA or P4/DHEAS were normalized to P4 only. Data are means ± SEM. **D-E**. representative traces for (b). **F**. The IC_50_ of E2 inhibition of the P4 response was calculated using the average current density at -80 mV for 8 cells recorded from HEK293 cells expressing human Kir7.1. **G**. Representative traces for (f). ML418 is a specific inhibitor for Kir7.1 (40).

Additionally, DHEA and P4 applied together showed a cumulative mode of action (Fig. 3B-D). While estrogen (E2) did not activate Kir7.1 upon direct administration (Fig. 3A), it was able to antagonize the P4 effect when added in combination. In fact, the IC_50_ of the E2 effect was 8.6 μM, indicating a competitive antagonist mode of action between E2 and P4, corroborating previous studies showing the inverse effects of progesterone and estrogen on uterine contractions (Fig. 3F-G and Suppl. Fig. 6); (1, 20).

While the steroid hormone progesterone (P4) sustains the developing fetus by maintaining uterine quiescence, another steroid hormone, estrogen (E2), transforms the myometrium into an active, contractile state. Progesterone has been widely recognized as a potent inhibitor of uterine contractions (26) with effects in micromolar concentrations, making it a principal factor in myometrium quiescence. However, the molecular mechanisms behind its effect on the myometrium are still unknown (3, 4-6, 27). Interestingly, the effects of both P4 and E2 on myometrial tissue are rapid, indicating a nongenomic mechanism of action, meaning that classical genomic steroid receptors that act via changes in gene expression are not the main regulatory units in this case. This exciting data open room for further extensive follow-up studies to explore whether other pregnancy-related hormones, such as estrogen, androgens, and dehydroepiandrosterone (DHEA), show any ability to regulate Kir7.1, perhaps via a competitive agonist/antagonist mode of action.

### Human Kir 7.1 responds to steroids used to treat pre-term labor

While medical intervention to induce contractions has been shown to be effective, prevention of uterine contractions, such as in the case of preterm births, has been proven more difficult without a complete understanding of uterine physiology. Preterm birth is defined as birth less than 37 weeks gestation and affects 10-12.5% of pregnancies in the US and accounts for 85% of perinatal morbidity and mortality (28). Annually, the societal cost of preterm births is $25.2 billion (29). Currently, vaginal administration of progesterone or intramuscular injections of an FDA-approved synthetic progestin, 17-alpha hydroxyprogesterone caproate (17-OHPC or Makena™), are the only approved treatments for the prevention of preterm labor (30,31).

Intramuscular administration of 250 mg 17-OHPC on a weekly basis starting at 16 to 20 weeks until delivery has been shown to significantly reduce the rate of preterm delivery (31). However, recently, larger studies have shown a less significant reduction in preterm labor (32). According to Sykes et al (33), this is due to a lack of understanding of the biological processes underlying preterm births and the heterogeneity of the population studied. Therefore, a greater understanding of the mechanisms underlying uterine contractions is essential for the prevention of preterm labor. By exploring 17OHPC action toward Kir7.1, we found that this compound not only specifically and potently activated Kir7.1 but also had a much stronger affinity towards Kir7.1, as demonstrated by a significantly slower washout property (Fig. 4A-B and Suppl. Fig. 7A-C). Our results show that Kir7.1 potentiation by 10 μM 17-OHPC was similar in strength to that of 10 μM P4; however, after drug withdrawal, the washout took five times longer with 10 μM 17-OHPC and was never fully washed out after exposure to a higher concentration of 50 μM 17-OHPC (Fig. 4A-B and Suppl. Fig.7A-B). This could explain why 17-OHPC has been shown to be effective with weekly injections, while vaginal progesterone needs to be administered daily (34).

**Figure 4.**
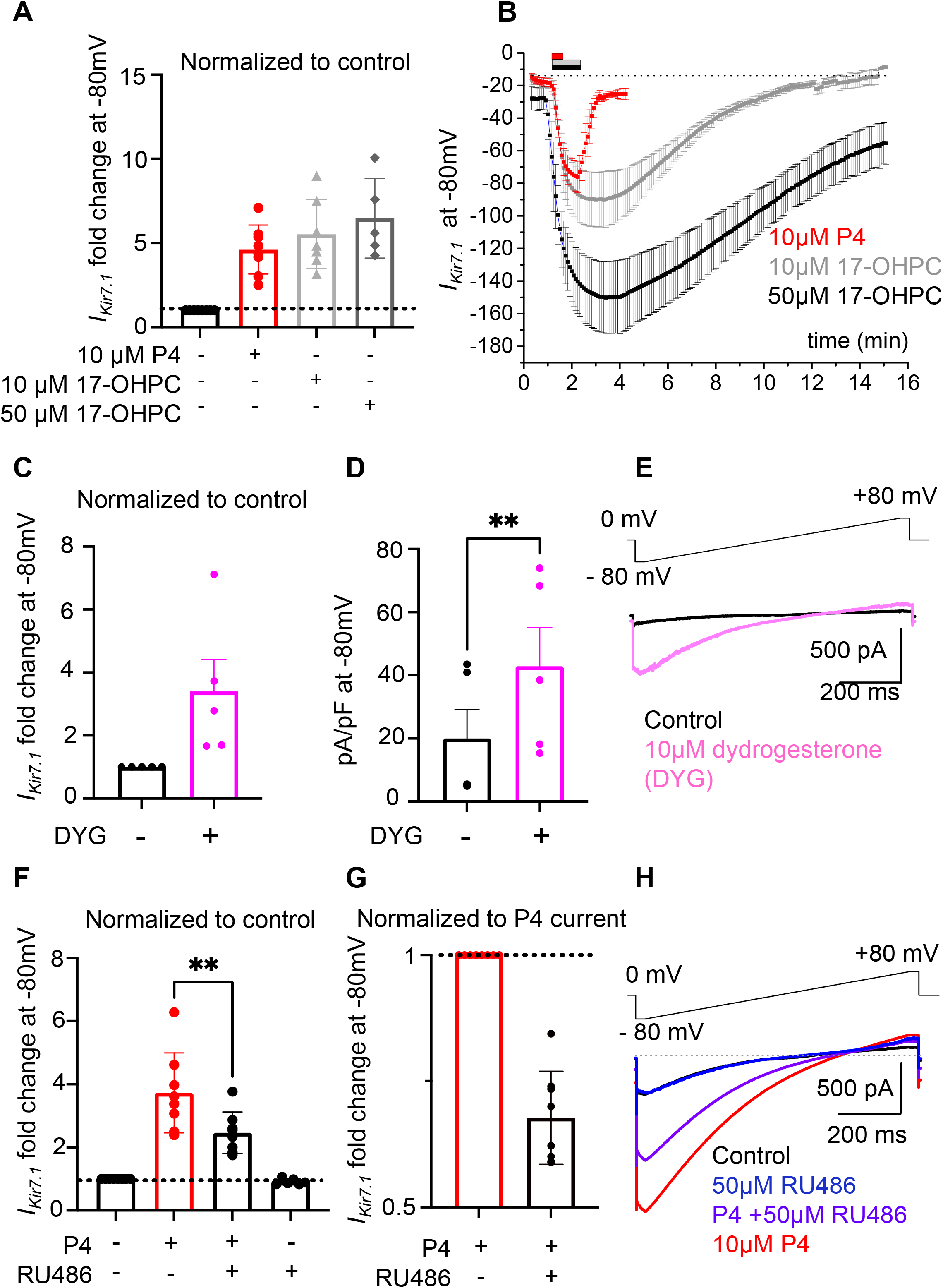
Regulation of human Kir7.1 by synthetic steroids. **A**. The average current fold increase of human Kir7.1 at -80 mV recorded from the recombinant channel expressed in HEK293 cells that were exposed to 10 µM progesterone (P4) or 10 µM and 50 µM 17-hydroxyprogesterone caproate (17-OHPC). **B**. Representative Kir7.1 current amplitudes obtained at -80 mV and plotted against time show a fast response to P4 and consequent fast washout, while the responses to 17OHPC are delayed and require a much longer washout time. The time of application of the compounds to the bath solution is indicated by the bars above. The responses were from HEK293 cells transfected with pIRES2-EGFP/K_ir_7.1. **C**. Graph depicting the fold change in the Kir7.1 currents as well as **D**. Kir7.1 current densities after exposure to 10 µM dydrogesterone (DYG) recorded from transfected HEK293 cells. **E**. Representative Kir7.1 traces recorded from HEK293-expressing human Kir7.1 cells stimulated with 10 µM DYG. **F**. Graph depicting the fold change in inward *I*_*Kir7.1*_ after exposure to 10 µM P4 recorded from transfected HEK293 cells either alone, in the presence of mifepristone (RU486) normalized to the control (f) or in response to P4 **(G). H**. Representative human Kir7.1 current amplitudes from HEK293 cells show that the P4 response could be partially antagonized by RU486. Statistical significance was calculated using the paired t-test, and the significance of changes is indicated as follows: **, *p* ≤ 0.01; n.s. stands for non-significant. Data are means ± SEM.

Another promising treatment for preterm labor currently in clinical trials is the synthetic progesterone dydrogesterone (Suppl. Fig. 7D). Dydrogesterone has been used since the 1960s worldwide in the treatment of threatened and recurrent miscarriages and was recently found to inhibit uterine contractions in a nongenomic manner (35-37). Here, we showed that Kir7.1 potentiation by dydrogesterone is similar to that of progesterone (Fig. 4C-E), further confirming the role of Kir7.1 in the fast, nongenomic regulation of uterine contractions. Dydrogesterone is also used in individuals to regulate the menstrual cycle and in the treatment of menstrual cramps.

### Kir 7.1 is inhibited by the abortifacient mifepristone

Towards the end of pregnancy, the quiescent uterus transforms into a highly contractile organ to allow for the expulsion of the baby. Endogenously, this phenotypic switch involves many factors, including functional progesterone withdrawal and increases in estrogen levels, oxytocin and prostaglandins. However, exogenous administration of 200-600 mg of the antiprogestin mifepristone (RU-486) alone induces contractions, thereby mimicking the onset of labor (38). RU-486 has also been shown to increase uterine sensitivity to prostaglandins and, when taken in succession, increases the efficacy of early pregnancy termination from 60% to 95% (39), making it a well-known abortifacient. As the first and only FDA-approved drug for early pregnancy termination, RU-486 has been administered to over 3.5 million Americans in the last two decades; however, the exact mechanisms of action have not been elucidated. The application of RU-486 to human Kir7.1 expressed in HEK293 cells revealed its antagonistic mode of action (Fig. 4F-H and Suppl. Fig. 7F) and show that RU-486 interferes with the P4 activation effect, uncovering the previously unknown mode of action for this compound.

Another risk factor for preterm labor is preeclampsia, a syndrome with clinical manifestations such as severe hypertension and edema. If left untreated, preeclampsia leads to fetal and maternal mortality. Placental insufficiency and impaired utero-placental fluid perfusion are thought to contribute to preeclampsia (1). Interestingly, the depletion of placental pericytes in the labyrinth is emerging as an important factor linked to preeclampsia development (1).

Progesterone and progestogens have been explored as a treatment against preeclampsia since the 20^th^ century, and recent studies indicated that dydrogesterone treatment showed statistically significant benefits (36). Placental pericytes have contractile properties and tightly enwrap placental capillaries and microvessels and form tight connections to the vasculature of the macrovessels (Suppl. Fig. 8 and Suppl. Fig. 2D). Hence, they are ideally positioned to control the blood exchange between maternal and fetal sites. Therefore, pericytial Kir7.1 upregulation by P4 is likely to keep the cells in a relaxed, hyperpolarized state and promote more effective fluid exchange. The absence of progesterone or pericytial insufficiency could alter this flow exchange by redistributing a large volume of placental blood back into maternal circulation, causing hypertension and edema while also producing fetal hypoxia. Further studies are needed to prove this hypothesis and evaluate the functional link between progesterone and Kir7.1 and their relation to pericyte insufficiency during preeclampsia.

Kir7.1 was previously identified as a crucial ion channel in maintaining membrane hyperpolarization during gestation, thereby promoting uterine quiescence (13). In this study, we determined the precise localization of Kir7.1 during pregnancy and revealed the physiological function of Kir7.1 in uterine quiescence through its regulation by both exogenous and endogenous steroids known to induce and/or inhibit uterine contractions. Moreover, taken together, this study shows the important role of Kir7.1 in the nongenomic regulation of endogenous and exogenous steroids during pregnancy, further deepening our understanding of uterine contractility and paving the way to develop selective pharmacological interventions to treat preterm labor.

## Figure Legends

**Supplemental Figure 1. Kir7.1 recording from nonpregnant and pregnant murine uteri. A**. Representative traces recorded from isolated myocytes from the nonpregnant uterus of 3-month-old mice show a lack of Kir7.1 activity and an absence of progesterone (P4). **B**. The average current fold change and the corresponding current densities (**C**) of mouse Kir7.1 at -80 mV recorded from isolated myocytes from nonpregnant uterus as in (A). **D**. Isolated myocytes from the nonpregnant uterus left to culture for 24 hours were identified by alpha-actin (cyan). **E**. Same day-isolated myocytes used for electrophysiological recording. **F**. Representative traces recorded from isolated myocytes from 15.5 dpc uteri of 3-month-old mice show variable Kir7.1 activity (populations A and B) and respond to P4 stimulation, which is further inhibited by the Kir7.1-specific antagonist VU590. **G**. Isolated myocytes from 15.5 dpc uterus in culture show both Kir7.1 (red) and alpha-actin (cyan) presence. The channel seems to be partially internalized. **H**. The average current fold change and the corresponding current densities (i) of mouse Kir7.1 at - 80 mV recorded from isolated myocytes from 15.5 dpc uterus as in (f-g). Data are means ± SEM.

**Supplemental Figure 2. Kir7.1 is asymmetrically expressed in murine placental pericytes. A-B**. Two representative immunostaining of isolated pericytes from 15.5 dpc uterus show both Kir7.1 (red) and alpha-actin (cyan) presence. The ion channel seems to be asymmetrically clustered around the cellular membrane. **C**. A cluster of isolated placental pericytes. **D**. The magnified portion of Fig. 2 shows placental pericytes forming tight connections with smooth muscles of the blood vessel. The border between two cells is indicated by yellow arrowheads.

**Supplemental Figure 3. Kir7.1 recording from murine placental pericytes. A-B**. The average current densities of mouse Kir7.1 at -80 mV recorded from two different populations of placental pericytes: population A and population B, as well as combined data (**C**). Lower panels: corresponding representative traces from 15.5 dpc pericytes show variable Kir7.1 activity (populations A and B) and respond to P4 stimulation, which is further inhibited by the Kir7.1-specific antagonist VU590. Data are means ± SEM.

**Supplemental Figure 4. Progesterone activates recombinant human Kir7.1 expressed in uterine smooth muscles as well as in HEK293 cells. A**. Representative traces recorded from HEK293 cells or human primary uterine smooth muscle cells (HUtSMCs). **B**. Transfection with a pIRES2-EGFP/K_ir_7.1 construct showed five-to ten-fold potentiation of the current after the addition of 10 µM progesterone (P4) to the bath. The cells transfected with the empty vector (upper panel) do not respond to P4 or display *I*_*Kir7.1*_. **C**. The average current fold change of human Kir7.1 at -80 mV recorded as described in (a). **D**. The average current fold change of human Kir7.1 at -80 mV recorded as described in (b). **E**. Human K_ir_7.1 (red) expressed in HEK293 cells transfected with the pIRES2-eGFP/K_ir_7.1 construct. **F**. Isolated HEK293 cells show clear membrane retention of Kir7.1. **G**. Western blotting detects both the mature glycosylated and shorter immature product of K_ir_7.1 seen in HEK293 cells transfected with pIRES2-EGFP/K_ir_7.1. The cells transfected with the empty vector did not express K_ir_7.1. Actin was used as a loading control. **H**. HUtSMCs transfected with recombinant pIRES2-eGFP/K_ir_7.1 (red). **I**. EC_50_ for P4 activation recorded from two different cell lines, HEK293 or HUtSMCs, transfected with human Kir7.1 was calculated using the average current density at -80 mV for n cells.

**Supplemental Figure 5. Progesterone activates recombinant Kir7.1. A**. The average current densities of human Kir7.1 at -80 mV recorded as described in Suppl. Fig 4a. **B**. The average current densities of human Kir7.1 at -80 mV recorded as described in Suppl. Fig 4b. **C**. EC_50_ for P4 activation recorded from HEK293 cells transfected with mouse Kir7.1 was calculated using the average current density at -80 mV for n cells.

**Supplemental Figure 6. Potentiation of human Kir7.1 by endogenous steroids**. Representative traces recorded from HEK293 cells transfected with a pIRES2-EGFP/K_ir_7.1 construct show different responses to the corresponding steroids. Right panels depict their chemical structure.

**Supplemental Figure 7. Effect of synthetic steroids on recombinant human Kir7.1**.

**A.** Representative washout traces recorded from HEK293 cells expressing human Kir7.1 following stimulation with 10 µM progesterone (P4), 10 µM hydroxyprogesterone caproate (17-OHPC), and 50 µM 17-OHPC. Sweep #1 in blue indicates the first sweep in Bath Solution following full potentiation by the steroids. The final sweep in red indicates the total washout. **B**. Graph depicting washout time in minutes for each steroid. **C-E**. Chemical structures of the exogenous steroids **F**. Graph depicting the potentiation by P4 and subsequent inhibition by RU486 at -80 mV.

**Supplemental Figure 8. Endothelial and smooth muscle expression in placental labyrinth**.

**A.** Horizontal cross-section through the labyrinth at 15.5 dpc; placental pericytes are visualized by their characteristic smooth muscle alpha-actin (cyan) staining, and capillaries are visualized by the endothelial marker CD31. **B**. Zoomed in section of (a) with pericytes stained with alpha-actin (pink arrow) connecting capillaries stained by CD31 (red arrow) **C**. Zoomed in section of (a) with individual smooth muscle alpha-actin (cyan) staining shows pericytes (pink arrow) and smooth muscle vasculature surrounding blood vessel (yellow arrow). **D-E**. single channel panels of (A)

## Acknowledgments

This work was supported by Pew Biomedical Scholars Award and Packer Wentz Endowment Will and funded in part by GCRLE grant from Global Consortium for Reproductive Longevity and Equality at the Buck Institute, made possible by the Bia-Echo Foundation, and NIH/NIA R03 grant (to PVL). The authors declare no competing financial interests. We thank Dr. Lin He lab and Elena Solonenko, M.Sc. for the help with cell preparations. The authors are grateful to Dr. Dimitra Gkika for critical reading of this manuscript.

## Author Contributions

Drs. Haoui and Lishko contributed to the conception and design of this research and wrote the manuscript. Dr. Haoui obtained most of all experimental data. All authors performed experiments, analyzed data, and reviewed and approved the final version of the manuscript.

