## Supplemental figures for "Kir7.1 is the physiological target for hormones and steroids that regulate uteroplacental function"

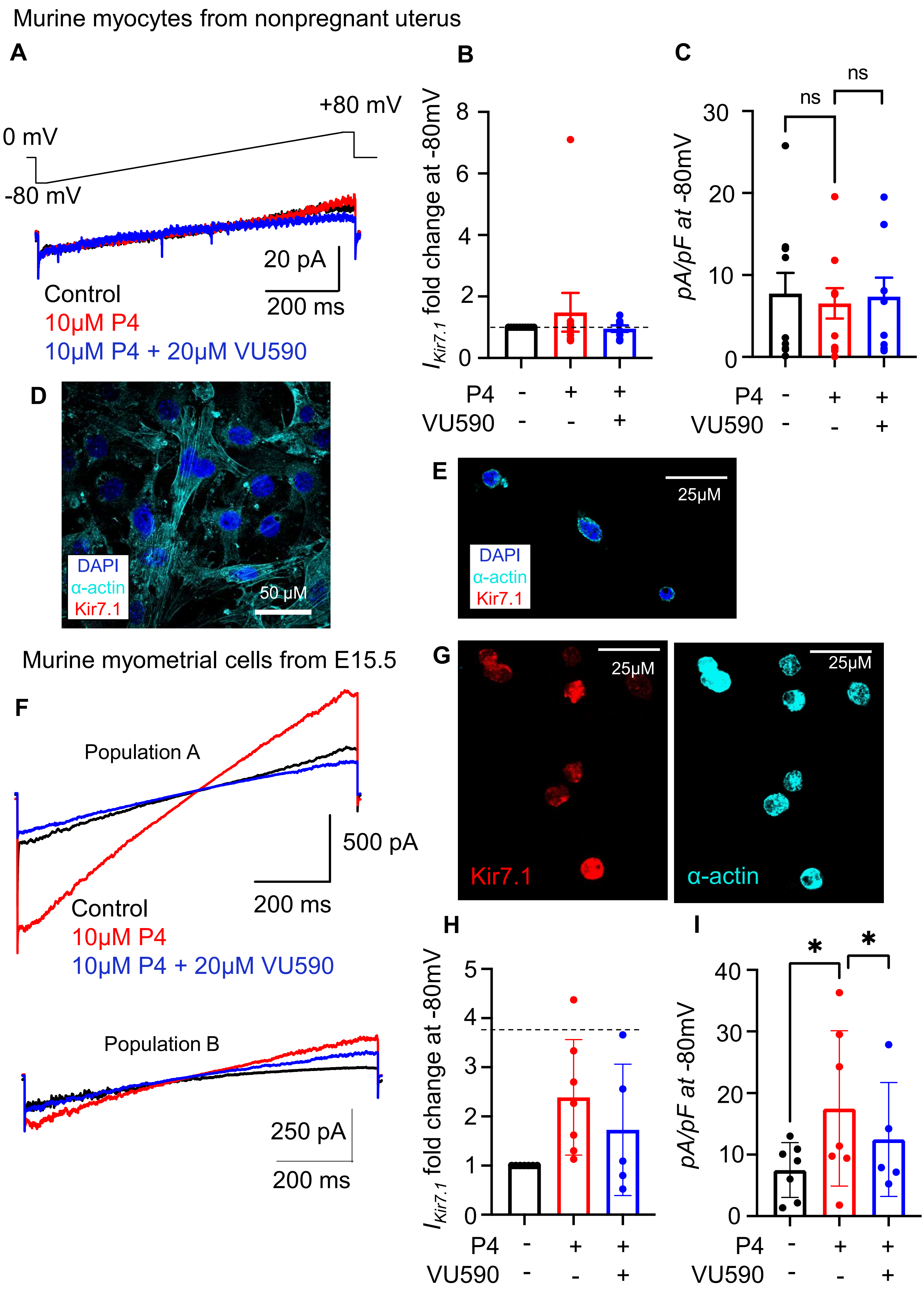


Supplementary Figure 1

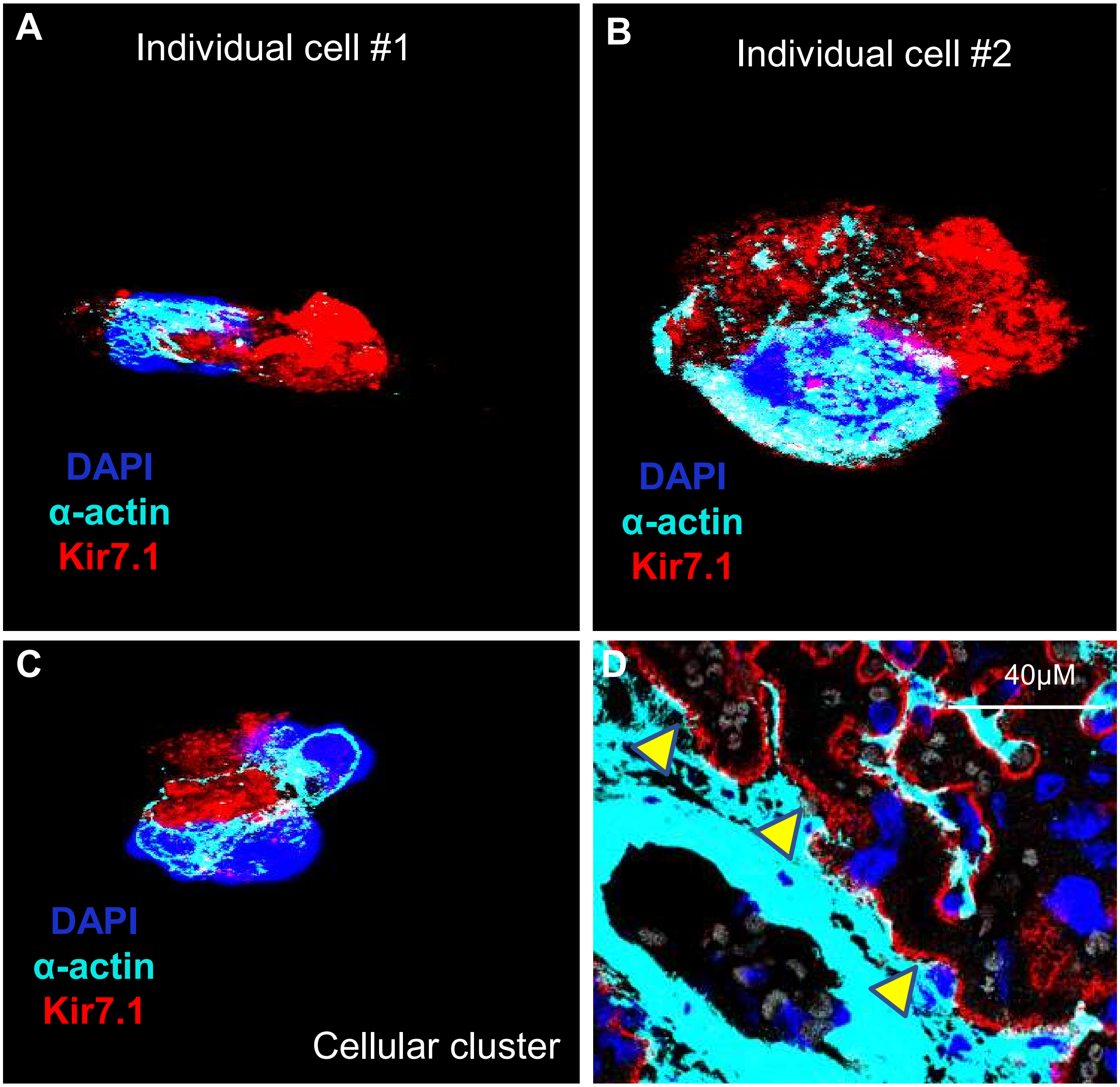


Supplementary Figure 2

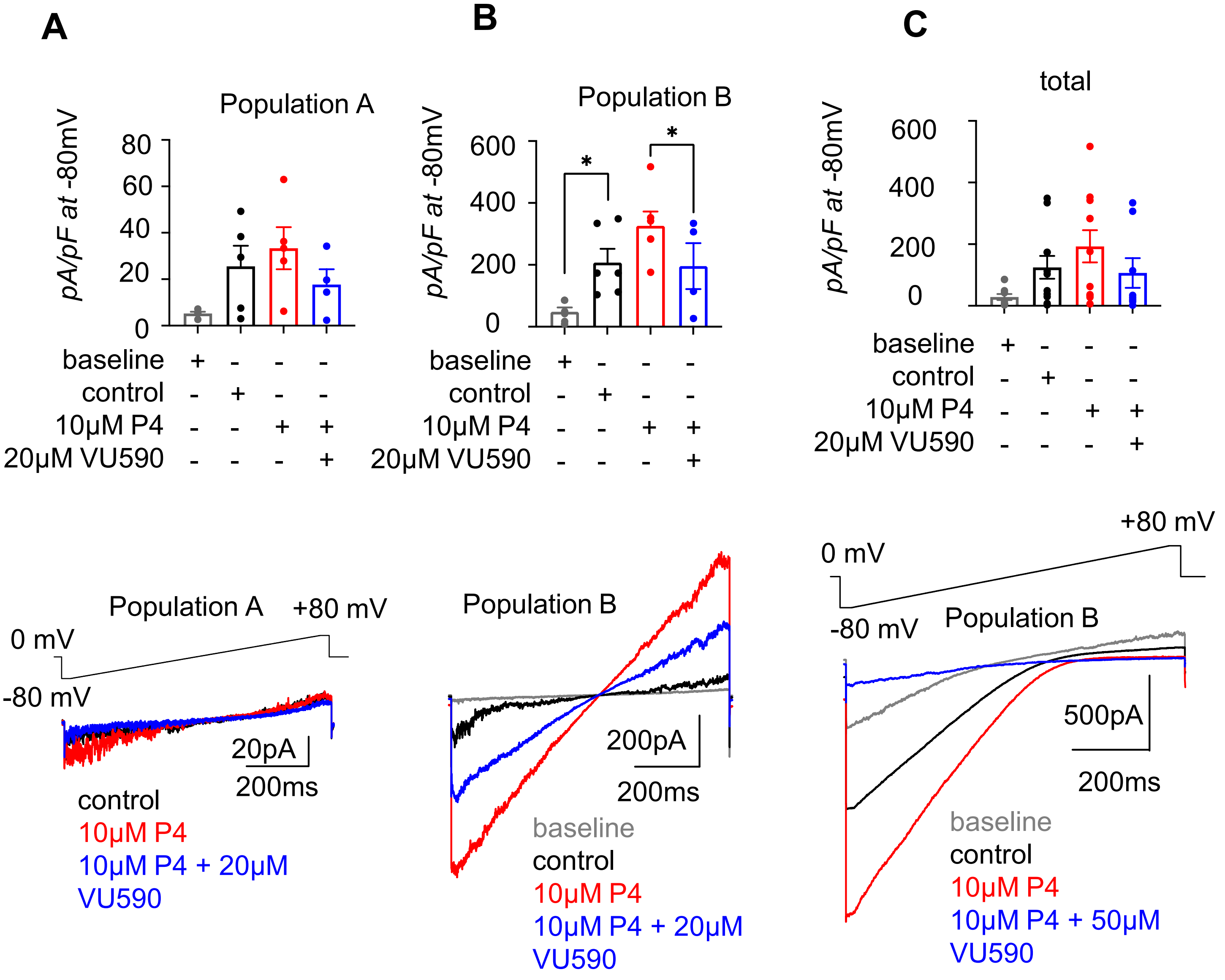


Supplementary Figure 3

Supplemental Figure 4

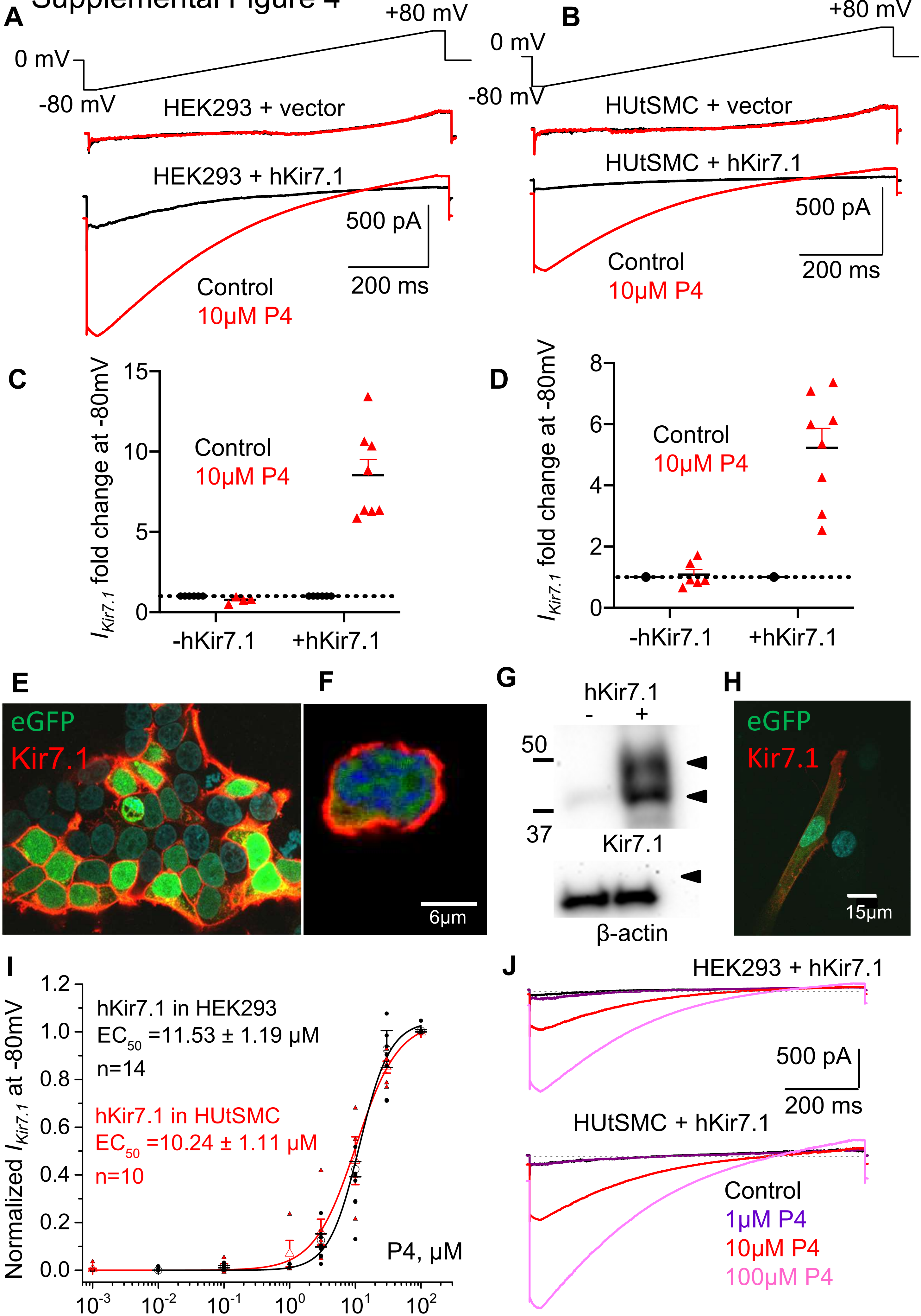

Supplementary Figure 5

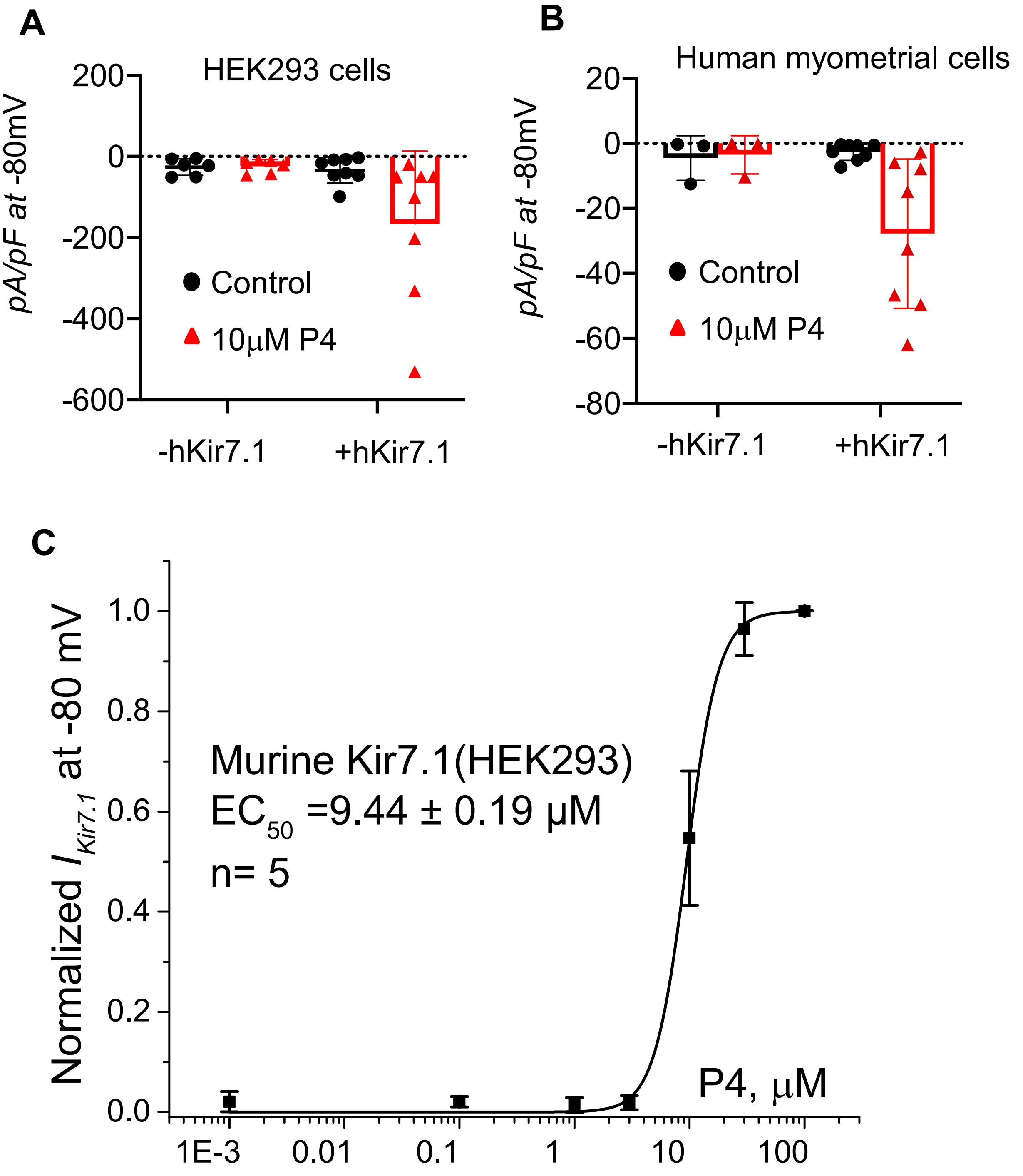

### Supplementary Figure 6

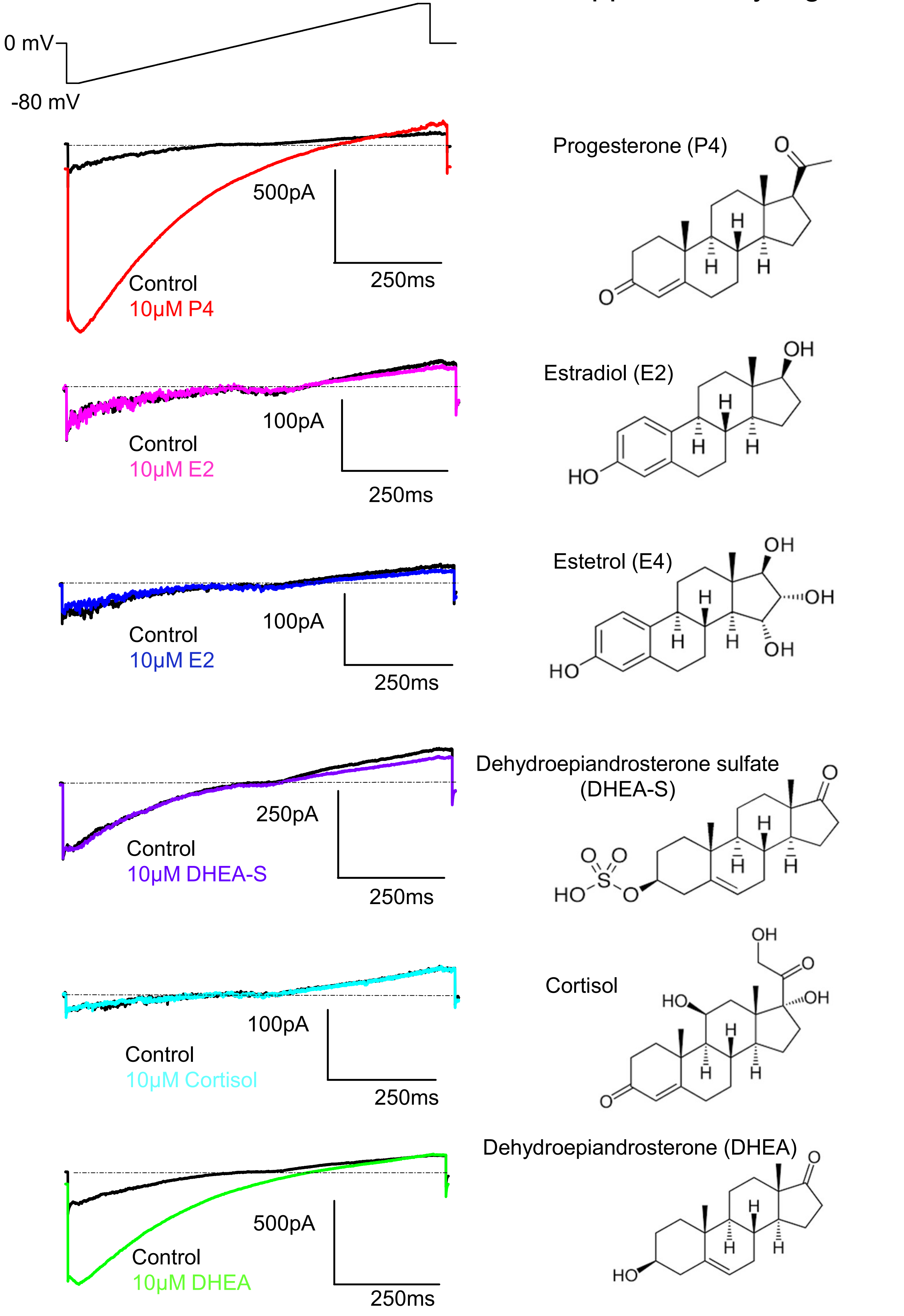

### Supplementary Figure 7

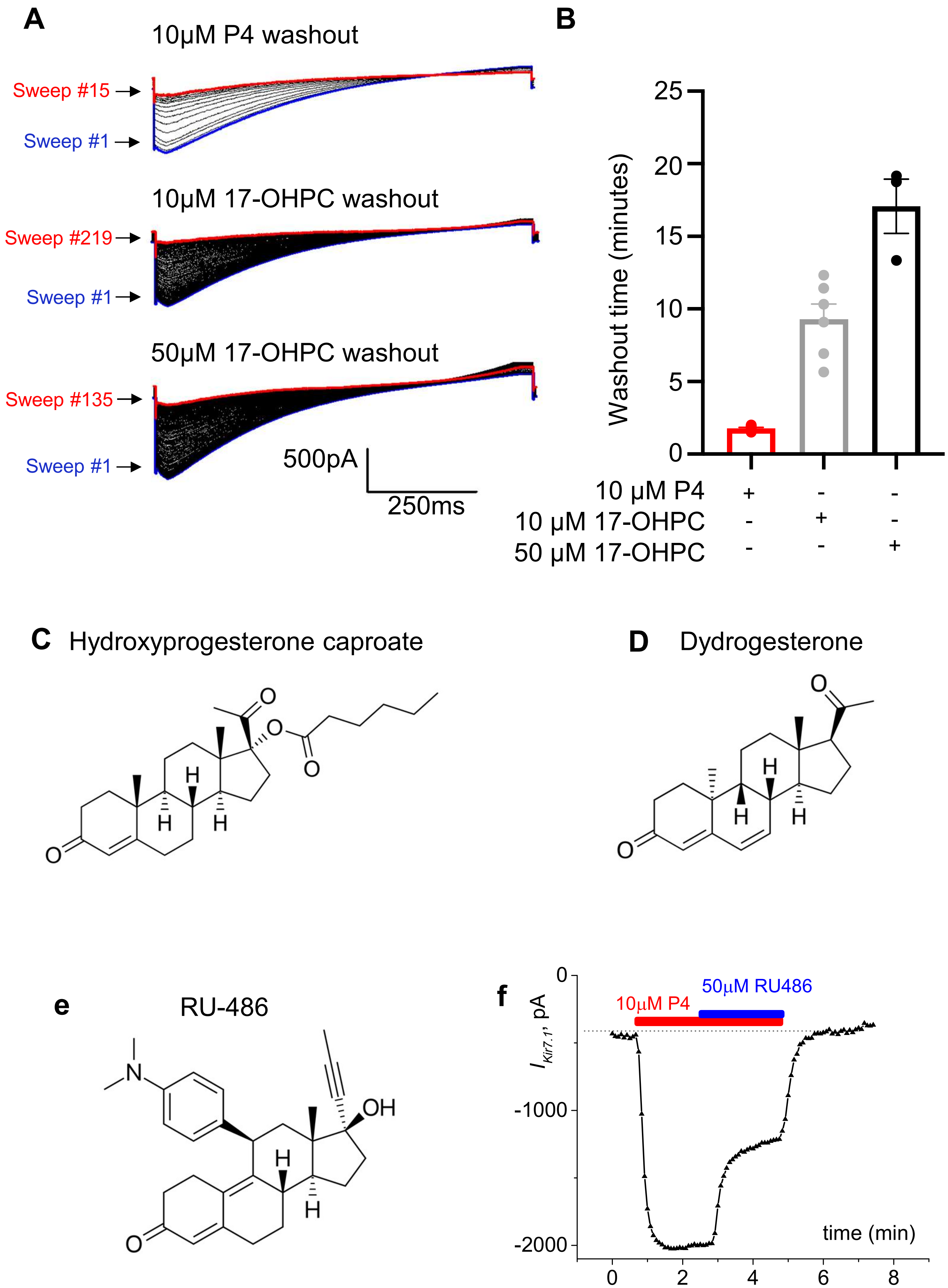

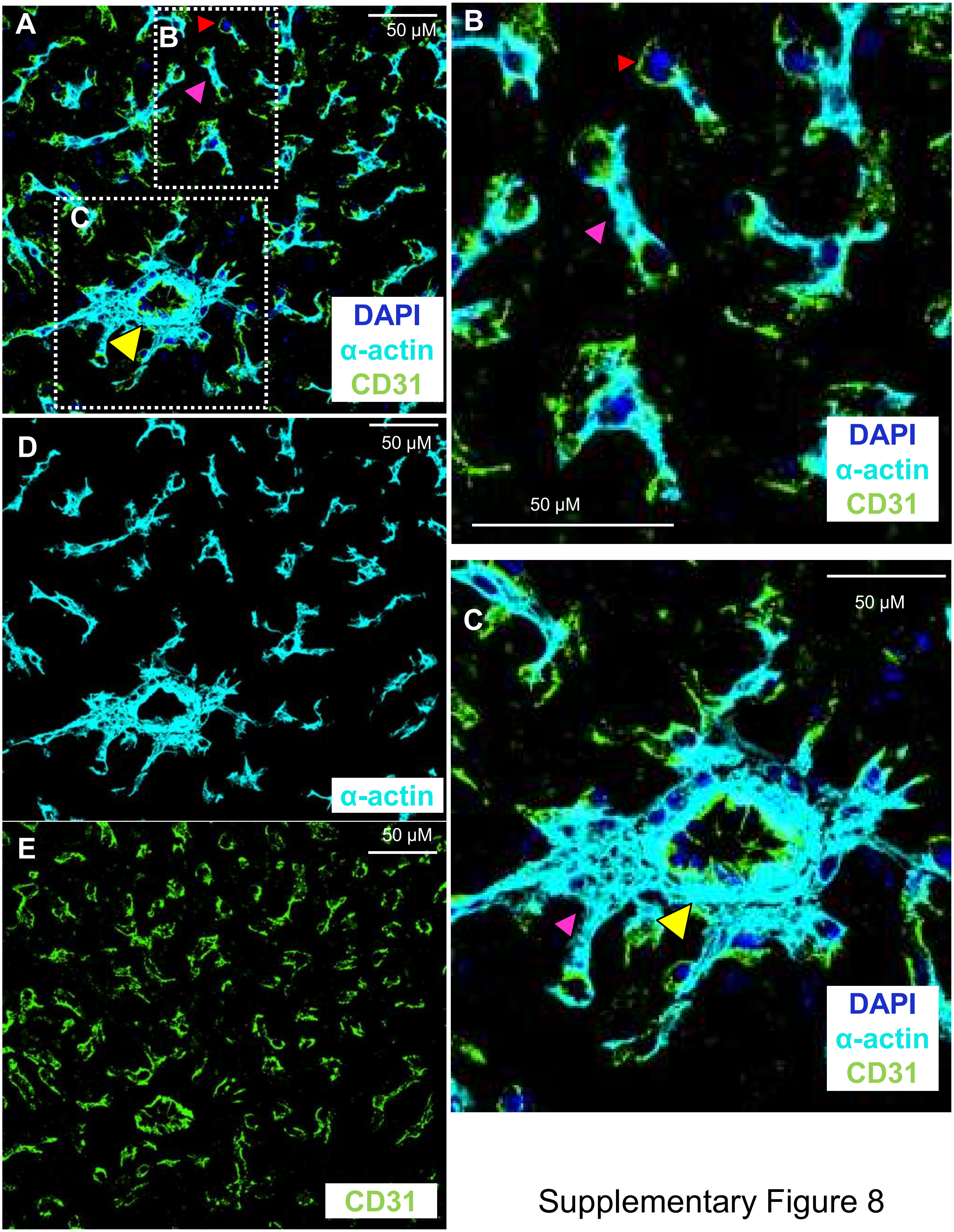


Supplementary Figure 8
